# Physiological Cross-Sectional Surfaces: A Method for Estimating Muscle Functional Capacity from 3D Digital Models of Fiber Architecture

**DOI:** 10.64898/2026.07.20.739018

**Authors:** Zhenhao Liu, Phoebe Duncombe, Vitaly Napadow, Geoffrey Handsfield

## Abstract

Physiological cross-sectional area (PCSA) is defined as the summed cross-sectional area of all muscle fibers contracting in parallel and at optimal length. PCSA is widely used, yet the conventional equations used to compute PCSA were developed from two-dimensional (2D) interpretations of muscle architecture and may not accurately represent muscle fiber cross-sections in the case of real three-dimensional (3D) geometries of muscles. Related measures of functional cross-sectional area (FCSA) and geometric cross-sectional area (GCSA) were also developed and interpreted with simplified 2D representations. Using realistic 3D muscle architectures derived from medical imaging, we sought to investigate whether conventional definitions of PCSA, FCSA, and GCSA represent the summed cross-sectional areas of all parallel muscle fibers, the fundamental definition of PCSA. We found that none of these measures consistently represented this definition. Thus, we introduce the physiological cross-sectional surface (PCSS), a curved surface within a muscle volume that is everywhere perpendicular to the local fiber direction. We estimated PCSS in 3D muscle surface meshes reconstructed from MRI data, using fiber orientations derived from Laplacian fiber reconstruction. PCSS-derived estimates were compared with PCSA, FCSA, and GCSA across six muscles representing five architectural classes. PCSS differed from all conventional measures, with the magnitude and direction of disagreement depending on muscle architecture. PCSS-to-PCSA ratios ranged from 0.771 to 1.399, while GCSA underestimated PCSS by up to a factor of 2.256 in bi- and multipennate muscles and overestimated it in muscles with more uniform fiber arrangements. PCSS demonstrated high geometric fidelity (perpendicularity>0.987) and robustness to fiber density across a tenfold range (coefficients of variation 0.39–3.95%). These findings indicate that conventional cross-sectional area measures do not consistently account for all fiber cross-sections in parallel within realistic 3D muscle geometries. PCSS provides a geometrically rigorous alternative that may improve estimation of functional muscle capacity from subject-specific imaging data.

## Introduction

Skeletal muscle serves as the primary actuator for human movement, generating the mechanical forces required for locomotion, joint stabilization, and postural control. Accurate characterization of muscle force-generating capacity is fundamental to musculoskeletal biomechanics, biomechanical modelling, and in assessments of pathological muscle. This functional capacity of a muscle depends on its composition and architectural properties, including fiber types, pennation angles, fiber length, and physiological cross-sectional area (PCSA) (1,2). Collectively, these architectural properties determine how contractile material is distributed and oriented within the muscle volume (3). Among these, PCSA is widely established as the primary geometric index of maximal isometric force-generating capacity, as it reflects the summed cross-sectional areas (CSA) of all fibers arranged in parallel when fibers are at optimal length, and thus represents the total contractile force a healthy muscle can produce (1,3,4).

Directly measuring every fiber’s CSA is experimentally impractical. Therefore, the most widely adopted estimation method computes PCSA as the ratio of total muscle volume (V) to optimal fiber length (L₀), where L₀ is the fiber length at which peak isometric force is produced (5). A trigonometric correction is applied to account for fiber obliquity to the external tendon, yielding the functional cross-sectional area (FCSA), computed as V multiplied by the cosine of the pennation angle (θ) divided by L₀ (6). Both formulations rest on the same simplifying assumption that the entire muscle can be represented as a flat cross-sectional plane, with pennation angle and fiber length each summarized by a single scalar value (6,7). However, a muscle’s force-generating capacity is shaped by the interplay of three-dimensional (3D) factors, including overall muscle size and shape, and the spatial distribution of fiber orientations throughout the volume, neither of which can be adequately captured by scalar summaries derived from a two-dimensional (2D) geometrically simplified muscle model (3,8). Pennation angle and fiber length are often assessed from 2D images, introducing potential errors in muscles with complex 3D fiber arrangements (9,10). These assumptions become increasingly untenable as architectural complexity increases, particularly in multipennate, fan-shaped, and circularly arranged muscles, where no single pennation angle or fiber length value can represent the full distribution of fiber geometry across the muscle volume.

The geometric cross-sectional area (GCSA) is an alternative measure defined as the maximum CSA of the muscle measured perpendicular to the line of fiber orientation (11). While GCSA is geometrically well-defined and does not require fiber length measurement, its correspondence to PCSA is architecture-dependent (1,11). In architecturally simple muscles, such as unipennate muscles, fibers can run nearly perpendicular to the GCSA plane, meaning this cross-section captures fiber cross-sections with minimal angular distortion, and GCSA may therefore approximate PCSA reasonably well. However, in bipennate and multipennate muscles, fibers approach the GCSA plane at oblique angles from multiple directions. Because GCSA is defined along a single reference orientation and does not account for fiber pennation, it captures only the projection of the fiber cross-sections onto that plane, and thus underestimates the functional force-generating capacity of the muscle (11). The accuracy of GCSA as a surrogate for PCSA thus cannot be assumed without knowledge of the internal fiber architecture of the specific muscle under investigation. Taken together, PCSA, FCSA, and GCSA each rely on scalar simplifications, whether through volumetric estimation, angular correction, or a fixed perpendicular cutting plane. Considering the various complexities of muscle morphology and fiber architecture in 3D, none of the areas described above robustly captures the CSA of all fibers arranged in parallel across muscle shapes and architectural arrangements.

Estimating PCSA by computing the CSA of each individual fiber perpendicular to its own local orientation would theoretically resolve the limitations of both conventional and geometric approaches. Rockenfeller et al. used a 2D simplified muscle model in which PCSA is defined as a projected cross-section plane that is perpendicular to a uniform fiber orientation, exceeding the boundaries of the muscle belly (11). This assumption cannot accommodate spatially heterogeneous pennation, fiber curvature, or continuous variation in fiber orientation across the muscle volume. Lee et al. extended PCSA estimation into 3D using Voronoi polygons constrained perpendicular to the local fiber orientation and then normalized to line-of-action direction using the cosine of pennation angle (12). This method demonstrates conceptual robustness over prior algebraic methods, though each polygon is still a flat 2D cross-section, and averaging cross-sections over fascicle length may underestimate PCSAs for muscles with high variance in cross-sectional areas over the fascicle length, e.g. fan-shaped muscles. In that study, the authors used volumetric fascicle trajectories traced throughout the entire muscle interior as input; these data were obtained from careful cadaveric dissection and digitization and are generally not widely available across research groups or from *in vivo* datasets. A simpler method of representing 3D fiber architecture in muscle is desirable, and would ideally incorporate a means of representing muscle fibers digitally from *in vivo* imaging data.

The Laplacian fiber reconstruction method is an attractive approach as it derives 3D fiber architecture directly from muscle geometry. By solving the Laplace equation over the interior of a muscle volume, the method generates a harmonic scalar field whose gradient defines a continuous fiber orientation field throughout the muscle (13). This method requires only a surface mesh and segmented origin and insertion regions as input, both obtainable from routine MRI, and produces anatomically plausible fiber trajectories without manual tracing or diffusion tensor imaging (DTI) acquisition, currently the most established approach for estimating 3D fiber orientation *in vivo*, validated against cadaveric microdissection in prior work (14). The original formulation by Choi and Blemker demonstrated that the Laplacian vector field produces physiologically realistic fiber trajectories in multiple muscles, showing overall agreement with published architectural measurements (7,13). Handsfield et al. subsequently validated the approach against DTI in eight adult medial gastrocnemius muscles, reporting no significant differences in pennation angle or fiber length between the two methods (15). These validations support the Laplacian approach as a reliable basis for fiber reconstruction in skeletal muscles. The resulting continuous fiber orientation field makes it possible to define a local cross-sectional surface perpendicular to the fiber direction at every interior point, providing the foundation for computing architecture-based PCSA throughout the 3D muscle volume.

To address the limitations of existing methods in capturing spatially varying 3D fiber architecture, we propose the physiological cross-sectional surface (PCSS), defined as the curved surface within the 3D muscle volume that is everywhere locally perpendicular to the fiber direction. Unlike the flat plane implicit in conventional PCSA, FCSA, and GCSA methods, the PCSS conforms to spatially varying fiber orientations. This study aims to develop a method for estimating PCSS from 3D muscle fiber architectural models constructed using the Laplacian fiber reconstruction approach, and to evaluate this method using muscle geometries derived from the Visible Human dataset and a whole-body MRI study (16,17). Here we compare PCSS-derived estimates against those produced by conventional PCSA, FCSA, and GCSA methods across six muscles spanning five distinct architecture types, namely unipennate, bipennate, multipennate, fan-shaped, and circular, selected to represent a robust range of architectural complexity encountered in the human musculoskeletal system.

## Materials and Methods

### Data acquisition

3D muscle geometries were obtained from two sources. The first was the publicly available Visible Human dataset by Andreassen et al., accessed via SimTK (https://simtk.org/projects/3d-vh-geometry), comprising segmentations derived from a single adult male cadaver (39 years old, 180 cm, 90 kg) (17). The second was derived from a whole-body MRI study (collected during development of the TARA program (16), in which a healthy adult male (54 years old, 170 cm, 70 kg) was scanned on a Siemens MAGNETOM Skyra (3T) using a 3D gradient echo protocol with an isotropic voxel size of 1.2 mm. For both datasets, muscles and aponeuroses were segmented using nn-Interactive and manually corrected on ITK-SNAP where necessary (18,19). Six muscles spanning five architecture types were included: unipennate (adductor brevis), bipennate (tibialis anterior, rectus femoris), multipennate (soleus), fan-shaped (pectoralis major), and circular (orbicularis oris). All segmentations were exported as stereolithography (STL) surface meshes.

### Fiber reconstruction

Fiber origin and insertion regions were identified by computing the overlap between the muscle surface mesh and either the bone or tendon surface mesh, depending on the anatomical attachment type of each muscle, followed by manual correction in MeshLab to ensure anatomical accuracy (20,21). Fiber trajectories were reconstructed using the Laplacian vector field simulation method, in which the Laplace equation was solved within the tetrahedral mesh of each muscle volume using the finite element method (13). Both steps were implemented using custom Python code. Neumann boundary conditions were applied at the origin (inflow flux, g_in = −1) and insertion (outflow flux, g_out scaled to balance total flux). The resulting scalar potential gradient was volume-weighted, smoothed, and traced as streamlines to reconstruct fiber paths. Reconstructed fiber architecture was validated against cadaveric dissection data from literature and further reviewed by an experienced anatomist to confirm reasonable accuracy of fiber tracts (7,22,23).

To inform future application to *in vivo* skeletal muscle DTI tractography, fiber trajectories were computed across ten fiber seeding densities in increments of 0.5 mm, with inter-fiber nearest-neighbor spacings ranging from 0.5 to 5.0 mm for most muscles and from 0.5 to 2.5 mm for the orbicularis oris because of its smaller volume. *In vivo* skeletal muscle DTI tractography conventionally uses voxel-based seeding, with seed points distributed throughout the muscle volume at specified spatial frequencies (24). Prior technical recommendations have suggested voxel volumes of approximately 20-30 mm³ for *in vivo* muscle DTI (25), correspond to isotropic voxel edge length of approximately 2.7-3.1 mm. Thus, the 0.5 mm spacing represents a substantially finer sampling density than the native voxel scale expected in *in vivo* DTI, whereas the 5.0 mm upper bound is coarser than this voxel-scale reference. Together, these fiber seeding densities test whether PCSS remains stable across a range that brackets plausible fiber-sampling densities for future voxel-based *in vivo* DTI applications. The numerical nature of this approach suggests that some effect of seeding density is expected. The stability across all ten fiber seeding density levels serves as an internal check of the method’s consistency.

### Conventional cross-sectional area estimation

For each muscle, PCSA was computed as V/L₀, where muscle volume V was derived from the segmented surface mesh and L₀ was the mean fiber length^1^ across all reconstructed fiber trajectories at each fiber density seeding level, computed as both the arc length along the trajectory and the straight-line distance between endpoints (Fig. 1A) (5). All fibers were assumed to operate at optimal length, as direct measurement of optimal fiber length *in vivo* is not currently feasible (11). FCSA was computed as V·cosθ/L₀, where θ is the pennation angle between each fiber’s mid-belly tangent and the muscle line of action, and cosθ was averaged across all fibers (6,26). The line of action was defined as the unit vector from the centroid of the origin region mesh to the centroid of the insertion region mesh, and cosθ was computed as the absolute dot product between each mid-belly tangent and this vector (Fig. 1C) (27,28). GCSA was computed by constructing a curved line of fiber orientation from the mean centroid path of all fiber trajectories, extending both ends to the muscle surface, resampling at 1 mm intervals, slicing the muscle mesh perpendicular to the local tangent at each point, and taking the maximum enclosed CSA (Fig. 1B) (11). All three measures were computed at each of the ten fiber density seeding levels to allow direct comparison with PCSS.

**Figure 1.**
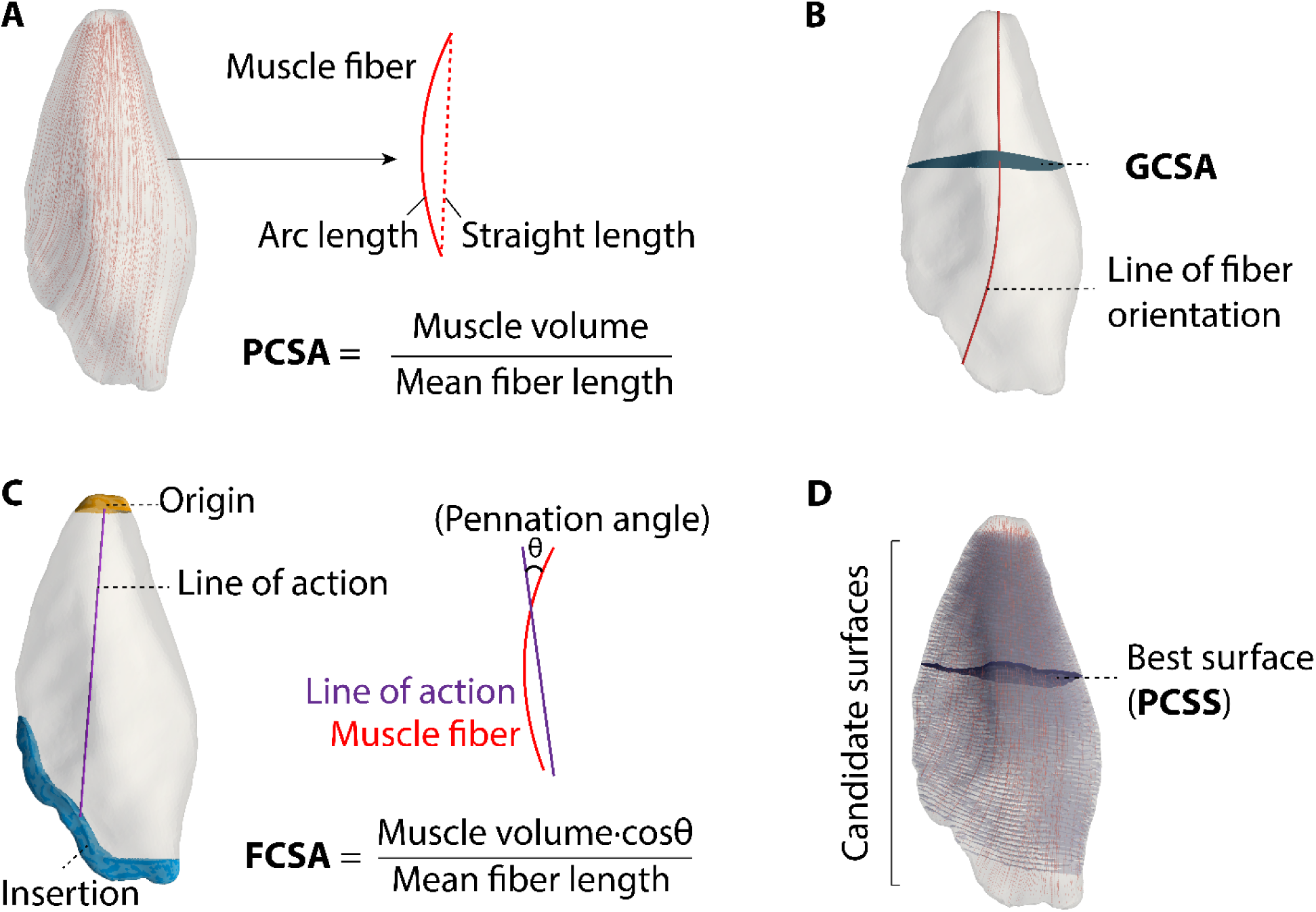
CSAs estimation methods. (A) PCSA = V/L₀, with L₀ computed as fiber arc length (solid) or straight-line length (dotted). (B) GCSA: maximum enclosed area within the muscle mesh that perpendicular to the line of fiber orientation (the mean centroid of all fiber trajectories). (C) FCSA = V·cosθ/L₀, where θ is the angle between each fiber’s mid-belly tangent and the muscle line of action (origin-insertion centroid vector). (D) PCSS: candidate isosurfaces (translucent) extracted from a potential field aligned with local fiber direction; the best-scoring candidate (dark surface) is selected by perpendicularity, fiber coverage, and surface balance.

### PCSS estimation

The PCSS of each muscle was estimated from the reconstructed fiber trajectories using custom Python code. A local fiber field direction was constructed by k-nearest-neighbor weighted interpolation of tangent vectors sampled along all fiber trajectories, and a scalar potential field was solved on a regular grid within the muscle volume by least-squares minimization, aligning the potential gradient with the local fiber field direction. Candidate perpendicular cross-sectional surfaces were extracted as isosurfaces of this potential field and clipped to the muscle boundary. Each candidate surface was evaluated on three geometric properties: perpendicularity (mean absolute dot product between surface normals and local fiber directions), fiber coverage (proportion of fibers intersecting the surface), and surface balance (proportion of non-intersecting fibers distributed symmetrically on either side of main surface). The highest-scoring candidate was selected as the main cross-sectional surface (Fig. 1D). Fibers not intersecting the main surface were spatially sub-grouped by their position relative to it, and an independent sub-surface was generated for each subgroup using the same potential field approach within the bounding box of that fiber subset. To ensure area estimates remained independent of fiber seeding density level, each fiber was assigned a representative CSA equal to the main surface area divided by the number of fibers crossing it. This per-fiber CSA defined a spatial influence radius used to trim regions of the sub-surface not intersected by any fibers, yielding a density-normalized sub-surface (Fig. 2). The total PCSS was computed as the sum of the main and all sub-surface areas, with perpendicularity retained as a quantitative index to assess geometric accuracy.

**Figure 2.**
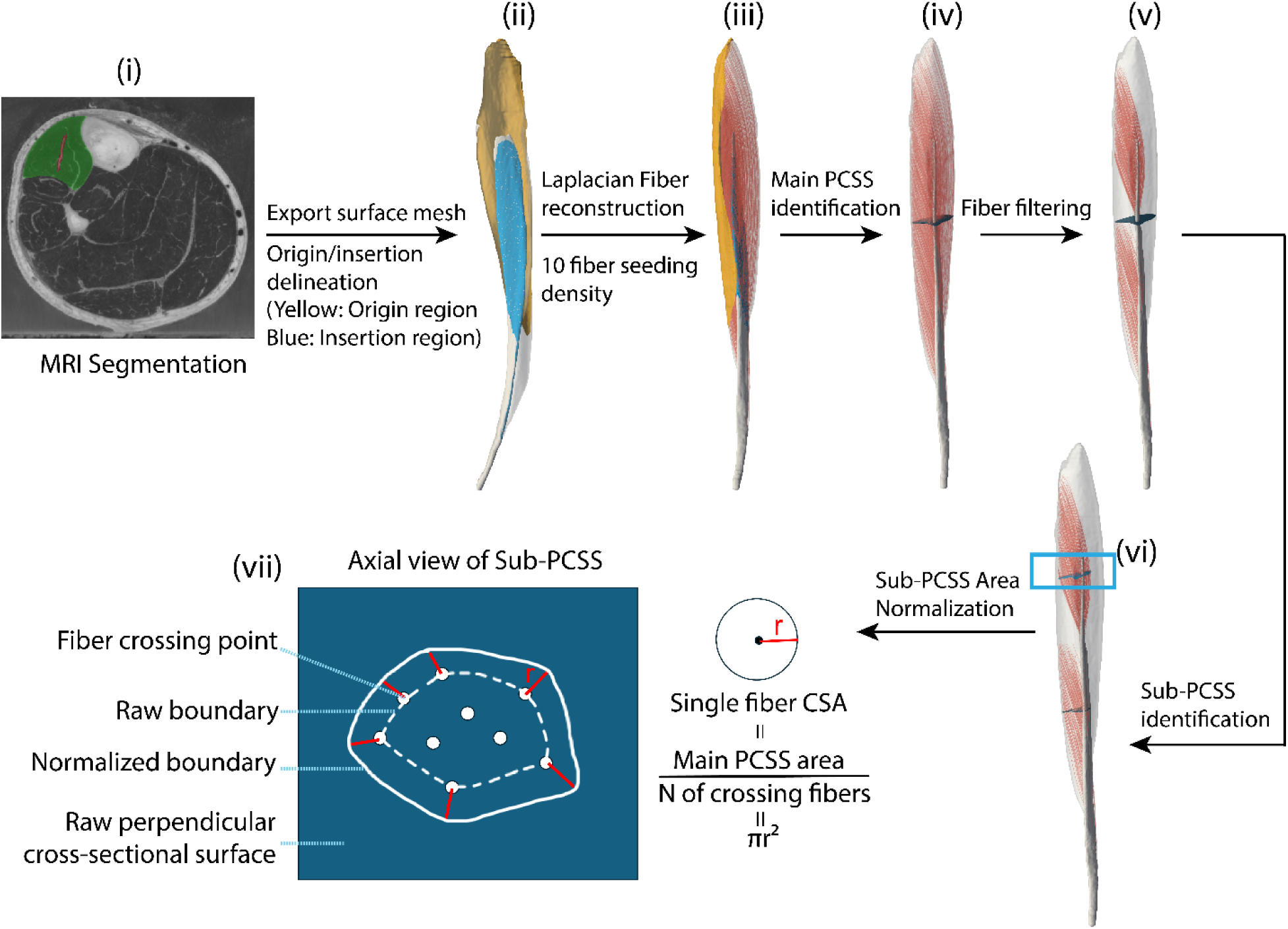
Overview of the automated PCSS estimation pipeline. (i) Muscle segmentation from MRI data, with origin and insertion regions delineated manually. (ii) Export of the surface mesh with origin (yellow) and insertion (blue) region labels, followed by Laplacian fiber reconstruction using a seeding density of 10 fibers. (iii) Main PCSS identification via a scalar potential field solved on a regular grid within the muscle volume, with candidate isosurfaces evaluated on perpendicularity, fiber coverage, and surface balance. (iv) The highest-scoring candidate is selected as the main PCSS. (v) Fiber filtering to identify fibers not intersecting the main surface, which are spatially sub-grouped for sub-PCSS generation. (vi) Sub-PCSS identification within the bounding box of each fiber subgroup using the same potential field approach. (vii) Axial view of a sub-PCSS illustrating the raw boundary (dashed line), normalized boundary (solid line) after area normalization, fiber crossing points (white dots), and the raw perpendicular cross-sectional surface. Each fiber is assigned a representative area (A) equal to the main surface area divided by the number of crossing fibers, and this per-fiber radius (r = √(A/π), red line) is used to trim sub-surface regions not covered by any fibers. The total PCSS is computed as the sum of the main and all sub-surface areas.

## Results

The PCSS produced estimates that differed from each of the conventional PCSA, FCSA, and GCSA methods, across all six muscles and all fiber density seeding levels tested (Fig. 3 and 4). The actual nearest-neighbor spacing achieved at each fiber density seeding level differed slightly from the target spacing due to the discrete nature of the mesh-based fiber seeding process, where the algorithm places fibers at the closest feasible positions within the muscle volume. In the adductor brevis, PCSA was 22-26% lower than PCSS (Fig. 4 A). In the bipennate muscles, tibialis anterior PCSA yielded values 14-32% higher than PCSS and rectus femoris PCSA yielded values 26-38% higher than PCSS (Fig. 4D, F). In the soleus, PCSA was similarly 9-21% higher than PCSS. In the fan-shaped pectoralis major and circular orbicularis oris, PCSA was 4-16% and 10-32% lower than PCSS, respectively (Fig. 4B, C, and E). FCSA followed the same directional pattern across all examined muscles, with values consistently lower than PCSA due to the pennation angle correction. GCSA showed a more variable pattern across architecture types. In the adductor brevis the ratio of GCSA to PCSS was near-equivalent (ratio = 1.02), while in the pectoralis major and orbicularis oris GCSA overestimated PCSS by factors of 1.11 and 1.12 respectively. By contrast, GCSA substantially underestimated PCSS in the rectus femoris (ratio = 0.44), soleus (ratio = 0.46), and tibialis anterior (ratio = 0.65). The magnitude of inter-method disagreement was largest in the bipennate muscles and soleus.

**Figure 3.**
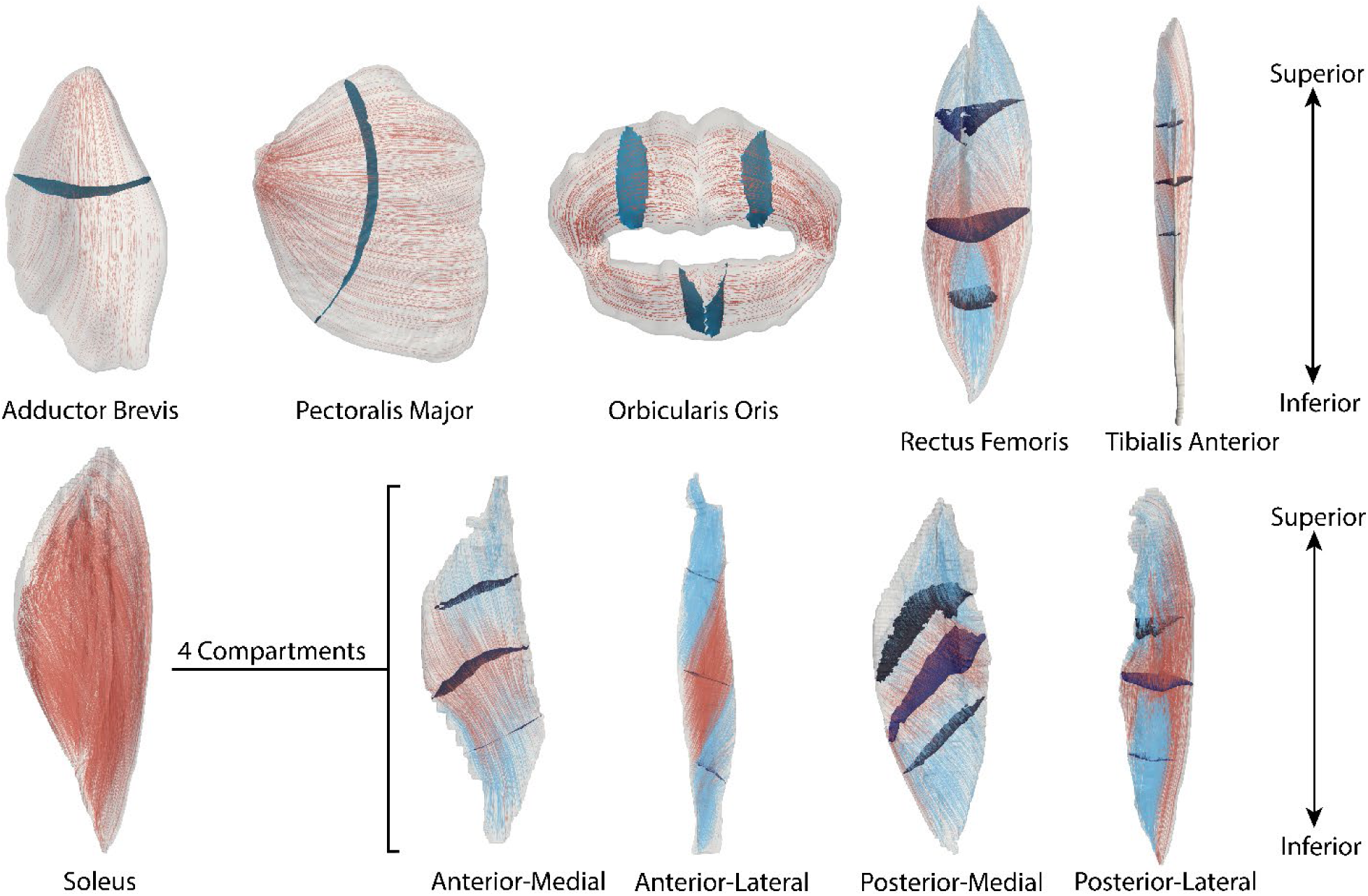
3D fiber architecture and PCSS (main surface and subsurface) for six muscles rendered on muscle meshes. The soleus is subdivided into four compartments (anterior-medial, anterior-lateral, posterior-medial, posterior-lateral), each shown individually with compartment-specific PCSS. Red fibers are covered by the main PCSS, while blue fibers are covered by the sub-PCSS.

**Figure 4.**
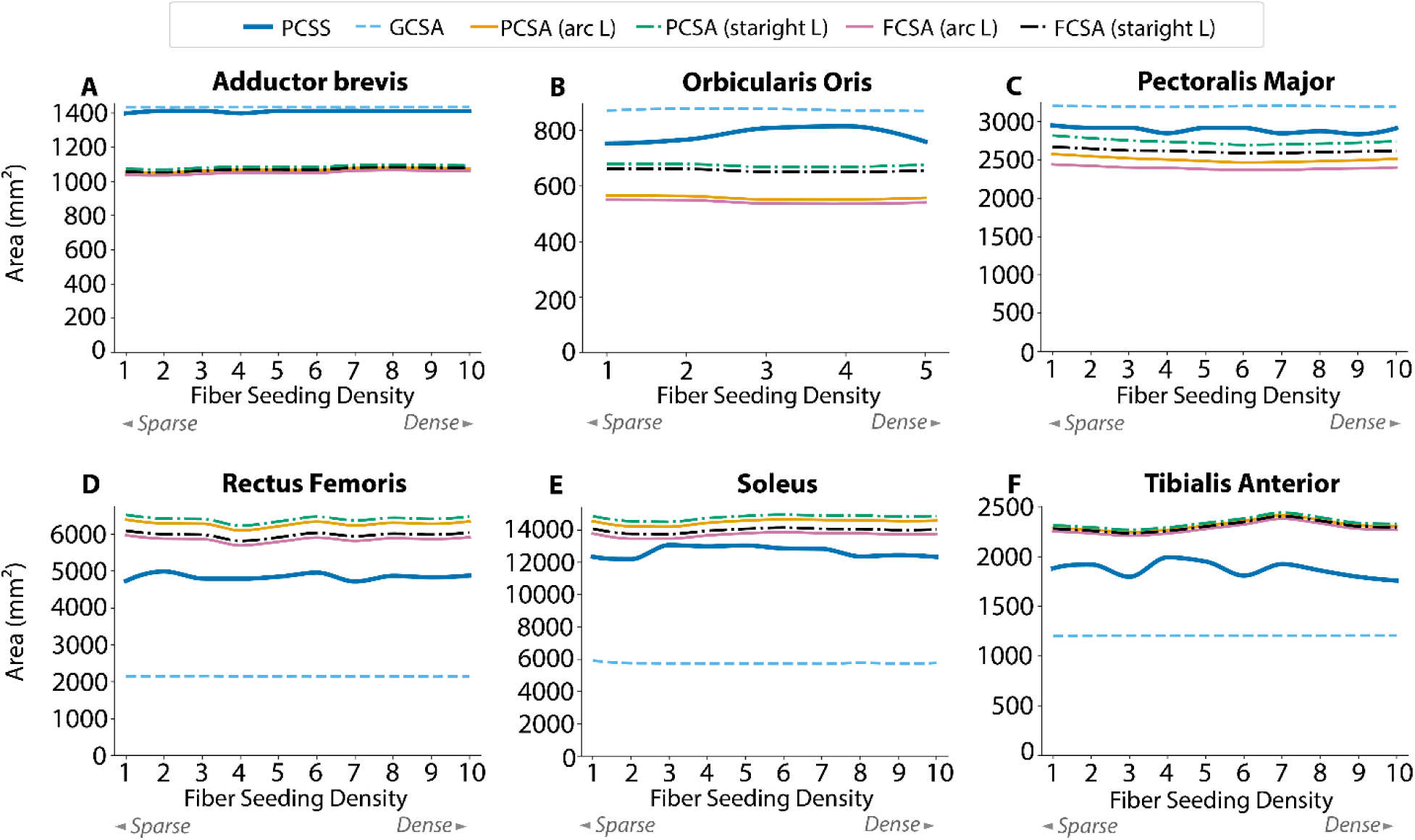
**Cross-sectional area & surface of skeletal muscle estimated by six methods across fiber seeding density levels** for (A) Adductor Brevis, (B) Orbicularis Oris, (C) Pectoralis Major, (D) Rectus Femoris, (E) Soleus, and (F) Tibialis Anterior. Line styles denote method type: PCSS (solid thick), GCSA max (dashed), arc length-based PCSA/FCSA (solid), and straight length-based PCSA/FCSA (dash-dot). Vertical separation between lines indicates inter-method differences in PCSA magnitude; flat trajectories indicate low sensitivity to fiber seeding density.

The perpendicularity of the PCSS was consistently high across all five architecture types and all fiber seeding density levels. Perpendicularity was expressed as mean ± standard deviation (SD): adductor brevis (0.9968 ± 0.0008), tibialis anterior (0.9904 ± 0.0021), rectus femoris (0.9876 ± 0.0045), soleus (0.9880 ± 0.0019), pectoralis major (0.9944 ± 0.0017), and orbicularis oris (0.9947 ± 0.0011). The highest perpendicularity values were observed in the adductor brevis and orbicularis oris, and the lowest in the soleus and rectus femoris, consistent with the greater spatial complexity of fiber orientation in these architectures.

PCSS showed coefficient of variation (CV) values of 0.39-3.95% across all examined muscles (adductor brevis 0.39%, tibialis anterior 3.95%, rectus femoris 1.73%, soleus 2.56%, pectoralis major 1.29%, orbicularis oris 3.32%), compared to 0.74-2.20% for PCSA and FCSA and 0.03-0.91% for GCSA. All methods showed generally stable estimates across the fiber seeding density range tested, with no systematic trend observed in any method. PCSS showed mild fluctuations in some muscles, particularly the tibialis anterior and soleus, while PCSA and FCSA remained relatively consistent across fiber seeding densities in most muscles. GCSA was essentially insensitive to fiber seeding density, showing near-zero CV and consistent values across all examined muscles and fiber seeding density levels (Fig. 5).

**Figure 5.**
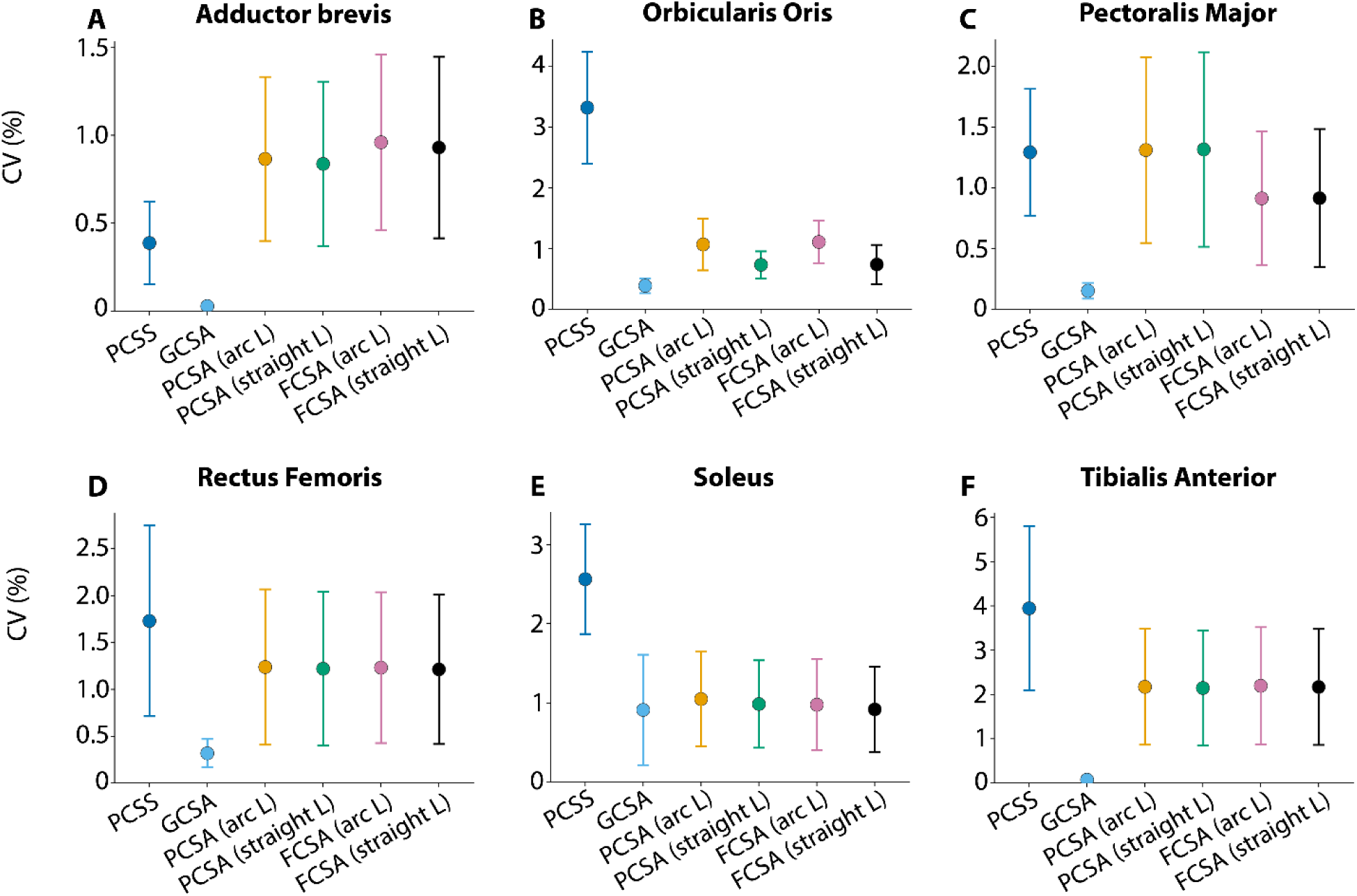
Sensitivity of six CSA methods to fiber seeding density. Coefficient of variation (CV = SD/mean × 100) of PCSA across ten seeding densities for (A) Adductor Brevis, (B) Orbicularis Oris, (C) Pectoralis Major, (D) Rectus Femoris, (E) Soleus, (F) Tibialis Anterior, across six methods (PCSS, GCSA, PCSA arc/straight L, FCSA arc/straight L). Error bars: SD of per-level absolute percent deviation from the mean (not a CI on CV). Lower CV = greater stability across seeding densities.

## Discussion

The present study introduced PCSS as a method for estimating muscle force-generating capacity that captures 3D fiber architecture and whole-muscle morphology, and compared it against conventional PCSA, FCSA, and GCSA across six muscles spanning five architecture types. The key findings are that PCSS produced different estimates from all three conventional methods, with the direction of disagreement differing across muscle architecture types, and that PCSS estimates were robust to fiber density seeding levels in all examined muscles. This robustness stems from PCSS’s ability to account for all muscle fiber CSA arranged in parallel directly from the muscle’s realistic 3D shape and fiber architecture, rather than reducing that architecture to a single length, angle, or centroid path as conventional methods do. By preserving this architectural information, PCSS provides a more accurate and reliable estimate of a muscle’s functional capacity.

The conventional PCSA formulation and its pennation-corrected variant FCSA have been widely adopted due to their computational simplicity and accessibility of input measurements. However, these formulations rest on assumptions that are difficult to satisfy in real muscle tissue. Fiber length in 3D space is not a single value but a spatially distributed quantity that varies across the muscle volume, and the pennation angle used in FCSA assumes a planar and uniform fiber orientation relative to a single line of action (9,11,29). Spatial variation in pennation angle within individual muscles has been documented across a range of architectural types, including muscles conventionally classified as unipennate (29). The present data illustrate that the direction and magnitude of disagreement between PCSS and PCSA differed between muscle architecture types, and in the case of the two bipennate muscles examined, between individual muscles sharing the same architectural classification (tibialis anterior: PCSS/PCSA = 0.808; rectus femoris: PCSS/PCSA = 0.771). This suggests that architectural classification alone does not fully predict the quantitative bias introduced by conventional methods, and that individual muscle geometry contributes additional variation even within an architecture type. Accordingly, the error introduced by the conventional formulation is not a fixed offset correctable by a single scaling factor, but varies with the specific 3D geometry of each muscle. PCSS avoids this problem by estimating CSA at the level of individual fibers using surfaces oriented perpendicular to the local fiber direction at each point, integrating both the morphological and architectural complexity of the muscle in 3D. The consistently high perpendicularity values indicate that the cutting planes were reliably oriented perpendicular to the local fiber direction across all architecture types and fiber density seeding levels.

The GCSA showed a pattern of disagreement with PCSS that was distinct from the PCSA and FCSA methods and consistent in direction with muscle architecture type. GCSA overestimated PCSS in the adductor brevis, pectoralis major, and orbicularis oris, while substantially underestimating in the rectus femoris, soleus, and tibialis anterior. This architecture-dependent pattern reflects the geometric relationship between fiber orientation and the measurement plane. When fibers traverse the GCSA plane at a large oblique angle, as in bipennate and multipennate muscles, the cut intersects fewer fibers than a PCSS would, resulting in underestimation; conversely, in muscles where fiber angles relative to the measurement plane are small, GCSA overestimates the PCSS. The present data systematically quantifies this bias across five distinct muscle architecture types, demonstrating that the direction and magnitude of the fiber discrepancy vary predictably with architectural type and that GCSA should be interpreted with caution particularly in bipennate and multipennate muscles where underestimation is most pronounced.

The low sensitivity of PCSS to fiber seeding density across a tenfold sampling range demonstrates that the method is robust to variation in fiber count, a property of direct relevance to *in vivo* application via DTI tractography. DTI can provide 3D muscle fiber orientation data in living subjects, enabling reconstruction of subject-specific fiber trajectories as input to the PCSS computation. This robustness is particularly relevant for clinical populations including older adults with sarcopenia, patients with neuromuscular disease, and individuals in post-surgical rehabilitation, where muscle architecture is altered and subject-specific 3D fiber geometry cannot be fully characterized by spatially limited conventional imaging methods. DTI tractography has been demonstrated to reconstruct altered muscle fiber architecture non-invasively *in vivo* across these populations, including aging (30), neuromuscular pathology (31), and post-stroke survivors (32), making it a natural input source for the PCSS computation in settings where conventional PCSA estimation from spatially limited measurements would be insufficient to capture subject-specific architectural heterogeneity.

PCSS and PCSA both estimate force-generating capacity along the fiber axis, but what actually drives joint movement is the force transmitted along the muscle-tendon unit axis, which is a distinction worth keeping clear. Like all CSA-based measures, PCSS does not directly represent tendon force. The transformation from fiber-direction force to tendon force additionally requires knowledge of the pennation angle (33). This distinction is relevant when interpreting PCSS values in the context of musculoskeletal modelling, where PCSS provides a more complete estimate of the total contractile force potential available at the fiber level, which can then be projected onto the tendon axis using the appropriate geometric relationships within the model.

There are several assumptions and limitations in this study. Each muscle architecture type was represented by a single muscle from a single individual, which limits the empirical generality of the quantitative comparisons reported here. The inter-individual variation in muscle geometry and fiber architecture may influence PCSS estimates in ways that cannot be assessed from the present data, and future work should examine PCSS across multiple subjects and specimens for each architecture type. Optimal fiber length was approximated by the anatomical fiber length derived from each reconstructed trajectory, as direct measurement of L₀ *in vivo* is not currently feasible (34). This approximation is consistent with prior PCSA studies employing MRI- or DTI-based fiber reconstruction and is justified by assumption that resting sarcomere lengths in many muscles are near-optimal under neutral joint positioning (34). No correction for sarcomere length deviation is applied, and in muscles that habitually operate away from optimal length the relationship between PCSS and isometric force capacity will be modified by the length-tension relationship (1,35). A further consideration arises in muscles containing intrafascicularly terminating fibers, such as the semitendinosus and sartorius, in which individual muscle fibers do not span the full fiber length but instead end and overlap within the fiber (36,37). In these muscles, intrafascicularly terminating fibers overlap within the fiber. Each fiber generates force proportional to its cross-sectional area, and since they run in parallel, their contributions sum across the fiber. Because terminating fibers do not insert into a tendon, they transmit their force laterally to the surrounding muscle tissue through shearing of the endomysium, rather than longitudinally through tension (38). A PCSS passing through this overlap region will intersect fibers from both terminating and spanning arrangements. Since terminating fibers transmit their force laterally through endomysial shear rather than directly to the tendon, it remains worth examining whether a cross-sectional measurement through this region fully and accurately captures the total force-generating capacity of the fiber. More broadly, PCSS estimates the mechanical potential of the contractile apparatus based on morphology and architecture alone, but actual force output *in vivo* is further modulated by fiber type, composition, activation level, and neural drive, none of which are captured by any CSA measure (1,2). Future work combining PCSS with finite element modelling and direct force measurement in experimental approaches will be necessary to establish the empirical quantitative relationship between PCSS and actual contractile force output across muscles.

## Conclusion

The present study introduced PCSS as a novel method for estimating muscle force-generating capacity that explicitly accounts for 3D muscle morphology and fiber architecture. Unlike conventional PCSA and FCSA formulations, which reduce the complex spatial distribution of fiber orientations, muscle shape, and size to scalar summary values, PCSS conforms to the locally varying fiber direction at every point within the muscle volume. PCSS estimates differed from conventional PCSA, FCSA, and GCSA across six muscles spanning five architecture types, with the direction of disagreement varying with muscle geometry and architecture. These indicate that 3D morphology and architecture together are essential determinants of force-generating capacity that conventional methods cannot adequately capture. Cutting plane perpendicularity consistently exceeded 0.987 across all examined muscles and fiber seeding density levels, and PCSS demonstrated greater robustness to fiber seeding density than conventional methods in all examined muscles. In the future, incorporation with *in vivo* DTI tractography and direct force measurements in animal models are desirable to further assess the prediction capabilities of PCSS for contractile force capacity, and to explore its potential for subject-specific clinical assessment of muscle function.

## Acknowledgment

The authors gratefully acknowledge the U.S. National Library of Medicine’s Visible Human Project for providing the open-source anatomical imaging data that served as the foundation for the musculoskeletal geometry used in this work. We also thank John Kirsch and the Martinos Imaging Center for their contributions to MRI data collection. This and VN were supported by the National Center for Complementary and Integrative Health (NCCIH), NIH (U24-AT012560). This work was supported by the Lampe Joint Department of Biomedical Engineering at UNC and NC State, and the UNC Department of Orthopaedics, whose resources and expertise were instrumental in the completion of this research.

## Footnotes

1 In this study, ‘fiber length’ refers to the distance between aponeuroses measured along the fiber’s orientation. Although ‘fascicle length’ is sometimes used in consideration of fibers sometimes terminating within a fascicle, the muscles examined here are not known to contain intrafascicularly terminating fibers.

